# Whole genome sequences and annotations of Japanese and French strains of *Heterosigma akashiwo*

**DOI:** 10.64898/2026.08.19.745619

**Authors:** Tsubasa Kondo, Mika Sakamoto, Mayuko Tokumaru, Yasuhiro Tanizawa, Yasukazu Nakamura, Atsushi Toyoda, Shoko Ueki

## Abstract

High-quality reference genomes provide an essential foundation for elucidating the molecular basis of organismal ecophysiology. Here, we sequenced and assembled chromosome-scale genomes of two *Heterosigma akashiwo* strains isolated from coastal waters of Japan and France. The assembly sizes were 1.18 Gb and 1.43 Gb for the Japanese and French strains, respectively. The scaffold N50 of the Japanese strain assembly was 66 Mb, whereas the one of the unscaffolded French strain assembly was 33 Mb. To our knowledge, these assemblies represent among the largest and most contiguous genome resources currently available for members of the Stramenopiles (Ochrophyta). Evidence-based gene prediction in the Japanese strain recovered approximately 90% of conserved stramenopile core genes, indicating a highly complete gene repertoire, and was complemented by extensive functional annotation. In the French strain, homology-based gene prediction recovered approximately 80% of conserved core genes. Comparative genome analysis revealed extensive synteny conservation between the two strains, although several putative duplication and translocation events were detected. These genomic resources provide a robust framework for investigating the molecular, cellular, and ecological mechanisms underlying the physiology, adaptation, and bloom-forming capacity of *H. akashiwo*.

## Introduction

Harmful algal blooms (HABs) are caused by noxious species and can exert detrimental effects on surrounding ecosystems as well as on local industries [1–5]. Elucidating the mechanisms underlying bloom formation and developing effective mitigation strategies are therefore critical for the management of HABs.

A range of environmental variables—including carbon dioxide concentration, nutrient enrichment (eutrophication), light intensity, temperature, and salinity—have been implicated in the initiation and persistence of HABs [1–3, 5–8]. Extensive field-based surveillance has characterized population dynamics of diverse HAB species and their correlations with environmental parameters, yielding substantial ecological insights. Complementary laboratory-based studies have further advanced understanding of the physiological responses of HAB-forming taxa under controlled conditions, providing important contributions to understand their ecophysiology. However, progress in elucidating the molecular mechanisms underlying HAB dynamics remains limited regardless advances of versatile cellular and molecular biology approaches. One of the major constraints is the insufficient availability of comprehensive genomic and genetic resources, which hampers in-depth investigation of gene regulation, functional genomics, and adaptive responses in HAB species.

*H. akashiwo* is a noxious raphidophyte that inhabits thalassic environments and frequently forms HABs during summer [1, 9, 10]. Although its toxicity has been debated for many years, the chemical identity of the toxic agent(s) has yet to be conclusively determined [1, 9–17]. Once considered primarily a temperate species, *H. akashiwo* has since been reported across both hemispheres, with a distribution spanning from Arctic to tropical regions, including the Pacific Rim, Oceania, and both the North and South Atlantic Oceans [4, 18–20].

The broad geographic distribution of this organism is likely attributable to its environmental adaptability, including tolerance to a wide range of temperatures [17, 21, 22], carbon dioxide concentrations [23, 24], and salinities[12, 21], as well as its ability to form cysts that enable survival under conditions unfavorable for vegetative growth [22, 25, 26]. In addition, resistance to turbulence [27] may facilitate long-distance passive dispersal via oceanic currents.

Moreover, the organism has been reported to phagocytose bacteria [28–30]. Consistent with these observations, our group recently demonstrated that *H. akashiwo* is capable of phagocytosing diverse bacterial species and utilizing them as a nutrient source under phosphate-depleted conditions. We further showed that *H. akashiwo* can proliferate to substantial levels when grazing on polyphosphate-accumulating bacteria, indicating that specific bacterial strains may promote its growth even under phosphate-limited conditions [31]. These findings highlight the importance of elucidating the metabolic flexibility of *H. akashiwo*, which encompasses both autotrophic and heterotrophic modes that may be differentially employed depending on environmental conditions.

Collectively, these studies suggest that *H. akashiwo* possesses a robust capacity to respond to a wide range of environmental challenges, likely supported by a versatile genetic repertoire and tightly regulated gene expression networks. The better availability of a high-quality genome sequence and comprehensive annotation is expected to substantially advance our understanding of the ecophysiology of *H. akashiwo*.

Here, we present the genome assembly and annotation of two *H. akashiwo* strains originating from Japan and France. Evaluation of assembly completeness suggests that the genomes, particularly that of the Japanese strain, achieve sub-chromosomal level resolution. These results establish a valuable genomic resource that will facilitate future studies of the molecular mechanisms underlying behavior and environmental adaptation of *H. akashiwo*.

## Materials and Methods

### H. akashiwo strains

*H. akashiwo* strain H93616 was originally obtained from the Japan Fisheries Research and Education Agency.

RCC1502 was obtained from the Roscoff Culture Collection (https://www.sb-roscoff.fr/en/roscoff-culture-collection). The strains were maintained in artificial seawater (ASW) supplemented with IMK medium (Fujifilm Wako Chemicals, Osaka, Japan) and antibiotics: penicillin (100 units/mL), streptomycin (100 µg/mL), ampicillin (100 µg/mL), and kanamycin (60 µg/mL). Cultures were maintained in a controlled-environment chamber under a 12 h light (100 µmol m⁻² s⁻¹) / 12 h dark photoperiod at 29°C for H93616 and 21°C for RCC1502. Both strains were cloned by limiting dilution prior to culture expansion and DNA extraction.

### DNA extraction and genome sequencing

High-molecular-weight genomic DNA was extracted from *H. akashiwo* H93616 and RCC1502 using a CTAB-based protocol, followed by purification with a Genomic-tip Kit (QIAGEN, Hilden, Germany) according to the manufacturer’s instructions [32]. The purified DNA was sheared to a target size of approximately 15–25 kb using a Megaruptor 3 system (Diagenode, Liège, Belgium). HiFi sequencing libraries were constructed using a SMRTbell Prep Kit 3.0 (Pacific Biosciences, Menlo Park, CA, USA) and size-selected for 10–50 kb fragments using a BluePippin system (Sage Science, Beverly, MA, USA). The libraries were sequenced on a Revio system using a Revio Polymerase Kit and a Revio Sequencing Plate (Pacific Biosciences, Menlo Park, CA, USA) with 30-h movie collections. Sequencing yielded 4.7 million HiFi reads (86.5 Gb) from one Revio SMRT Cell for H93616 and 7.5 million HiFi reads (106.5 Gb) from two Revio SMRT Cells for RCC1502.

For short-read sequencing, genomic DNA was fragmented to an average size of approximately 500 bp using a Focused-ultrasonicator M220 (Covaris, Woburn, MA, USA). A paired-end libraries were prepared using a TruSeq DNA PCR-Free Library Prep Kit (Illumina, San Diego, CA, USA) for H93616 and an Elevate Mechanical Library Prep Kit (Element Biosciences, San Diego, CA, USA) for RCC1502. The libraries were sequenced on a NovaSeq 6000 system (Illumina, San Diego, CA, USA) and an AVITI24 system (Element Biosciences, San Diego, CA, USA), respectively, to generate 2 x 150 bp paired-end reads. Library concentration and quality were assessed using a Qubit 4 Fluorometer (Thermo Fisher Scientific, Waltham, MA, USA), a 2100 Bioanalyzer system (Agilent Technologies, Santa Clara, CA, USA), and a 7900HT Fast Real-Time PCR System (Thermo Fisher Scientific).

### Analyses of *H. akashiwo* strains H93616 and RCC1502 genome charact**e**ristics

Genome characteristics were estimated based on Illumina paired-end reads. Illumina reads were subjected to *k*-mer counting using KMC (v3.2.4) [33] with a k-mer size of 21. The *k*-mer frequency distribution was analyzed using GenomeScope 1.0, which models sequencing errors, heterozygosity, and repetitive elements. Genome size, heterozygosity rate, and repeat fraction were inferred from the best-fit model[34].

### Genome assembly and phasing for H93616

The workflow of *H. akashiwo* H93616 genome assembly and scaffolding is presented in Fig1.

For *de novo H. akashiwo* strain H93616 genome assembly, PacBio HiFi reads and Hi-C sequencing data were utilized. Hi-C reads were preprocessed using fastp (v0.24.0, https://github.com/opengene/fastp, [35–37] ) to trim five nucleotides from the 5′ termini. The HiFi reads were then assembled using hifiasm (v0.25.0) to generate an initial assembly graph [38–40].

Subsequently, Hi-C data were integrated to enable haplotype phasing using the Hi-C-assisted phasing module implemented in hifiasm, the assembly was classified to a primary assembly and alternative contigs. The assembled sequences were extracted from the assembly graph in FASTA format and subsequently used for downstream analyses.

To remove potential contaminants, the assembled genome was screened using FCS-GX (v0.5.4-8-g3c7c426) [41], and contigs of bacterial origin were excluded. To obtain nuclear genome assembly, contigs representing *H. akashiwo* organelle genomes (GenBank accession No. KU561547 for mtDNA and LC269923 for cpDNA) and rRNA gene clusters were identified and removed using cleanup_short_contigs pipeline (https://github.com/nigyta/cleanup_short_contigs). The dataset was analyzed by SeqKit (v2.3.0) [42]to obtain statistic parameters for the assembly and the completeness of the assembly was evaluated by BUSCO v6 package based on Stramenopiles_odb12 dataset [43, 44].

### Hi-C scaffolding for H93616

Hi-C reads were mapped to the obtained contigs according to Arima-HiC Mapping Pipeline (https://github.com/ArimaGenomics/mapping_pipeline). Hi-C scaffolding was conducted by using cleaned contigs using YaHS (v1.2.2) [45]. The assembly structure was confirmed by visualizing Hi-C contact map using JuiceBox (v2.15) [46], and the dataset resulted from scaffolding was evaluated by SeqKit (v2.3.0) for assembly statistics and by BUSCO v6 for completeness.

The assembly was analyzed by using TIDK (v 0.2.65) to search for telomeres [47]. Positions of the identified telomeres were visualized with teloviz (https://github.com/mtok-omu/teloviz).

### Identification of organelle genomes for H93616

mtDNA and cpDNA were identified from the assembled contigs based on the sequence similarity to the previously published references (GenBank accession No. KU561547 and LC269923). For mtDNA, candidate contigs were aligned to the reference genome using MAFFT (v7.526), and the contig with the highest alignment identity and coverage was selected. Similarly, cpDNA candidate contigs were identified and aligned to the reference sequences using MAFFT [48, 49] and D-GENIES (https://dgenies.toulouse.inra.fr) [50] and both forward and reverse-complement orientations were evaluated to determine the optimal alignment. Because the organelle genomes are circular, overlapping regions at the end of these contigs are identified on sequence redundancy and removed to obtain the complete circularized genome sequences.

### RCC1502 genome assembly

The workflow of *H. akashiwo* RCC1502 genome assembly is presented in Fig1.

The HiFi reads were assembled using hifiasm (v0.25.0) to generate primary contigs. The assembly was screened using FCS-GX (v0.5.4-8-g3c7c426), and contigs of bacterial origin were excluded. The contigs representing *H. akashiwo* organelle genomes (GenBank accession No. KU561547 for mtDNA and LC269923 for cpDNA) and rRNA gene clusters were identified and removed using cleanup_short_contigs pipeline to obtain draft assembly. The assembly was analyzed using TIDK (v 0.2.65) to search for telomeres. The dataset produced from each step was analyzed by SeqKit (v2.3.0) to obtain statistic parameters for the assembly and completeness of the draft sequences was evaluated by BUSCO v6 package based on Stramenopiles_odb12 dataset.

### Gene prediction and functional annotation

Structural annotation of the H93616 and RCC1502 genome assembly was performed using both a deep learning–based approach (Helixer) [51, 52], and an evidence-driven annotation pipeline (BRAKER4) [53–58]. Helixer was executed on the web server (https://www.plabipd.de/helixer_main.html) using both land plant and vertebrate lineage models. The resulting gene models from the two lineage-specific runs were merged using AGAT (v1.7.0, https://github.com/NBISweden/AGAT) to obtain annotation based on combined model.

For evidence-based annotation of H93616, BRAKER4 was run in ETP mode, incorporating RNA-seq datasets from previous studies (DDBJ SRA accession No. DRR198893–DRR198900, DRR198909, and DRR198910) as transcript evidence, along with the OrthoDB Eukaryota protein dataset (Eukaryota.fa) as protein homology support. Liftoff(v1.6.3, https://github.com/agshumate/Liftoff) was adopted to structurally annotate RCC1502 genome using BRAKER4-based annotation of H93616 as a referance [59]. The resulting annotation datasets from different prediction tools were evaluated for completeness using BUSCO v6 in protein mode.

For organelle genome, the annotation of the references, KU561547 and LC269923, were transferred to the novel mtDNA and cpDNA sequences by LiftOff (v1.6.3) package [59]. The organelle genome annotations were integrated to the nuclear genome annotation to complete the whole genome sequence annotation.

Functional annotation was performed using the representative amino acid sequences of the predicted loci obtained from the BRAKER4 annotation (longest.aa.fa). KEGG Orthology (KO) identifiers were assigned using the KEGG Automatic Annotation Server (KAAS; https://www.genome.jp/kegg/kaas/) [60] with GHOSTZ and the bidirectional best-hit (BBH) method. Protein domain prediction, Gene Ontology (GO) term assignment, and other functional signature identification were conducted using InterProScan5[61], integrating multiple protein signature databases to generate comprehensive functional annotations.

### Synteny Analyses

Synteny comparison between H93616 and RCC1502 genome assembly datasets were conducted by GENESPACE v1.3.1[62].

## Results

### Initial characterization of the *H. akashiwo* genome

Statistics for the sequence reads obtained for this study are presented in Table 1. To estimate the genome size and heterozygosity of *H. akashiwo* H93616, 154.1 Gb Illumina reads were used for k-mer analysis by GenomeScope (Fig 2A). The estimated genome size of the strain H93616 was calculated to be 1.1Gb, and the estimated heterozygosity rate was approximately 2.14%. For RCC1502, 131.1 Gb Illumina reads were analyzed by the same procedure. The estimated genome size and heterozygosity for the strain is 1.21Gbp, and 2.45%, respectively (Fig 2B).

**Fig. 1.**
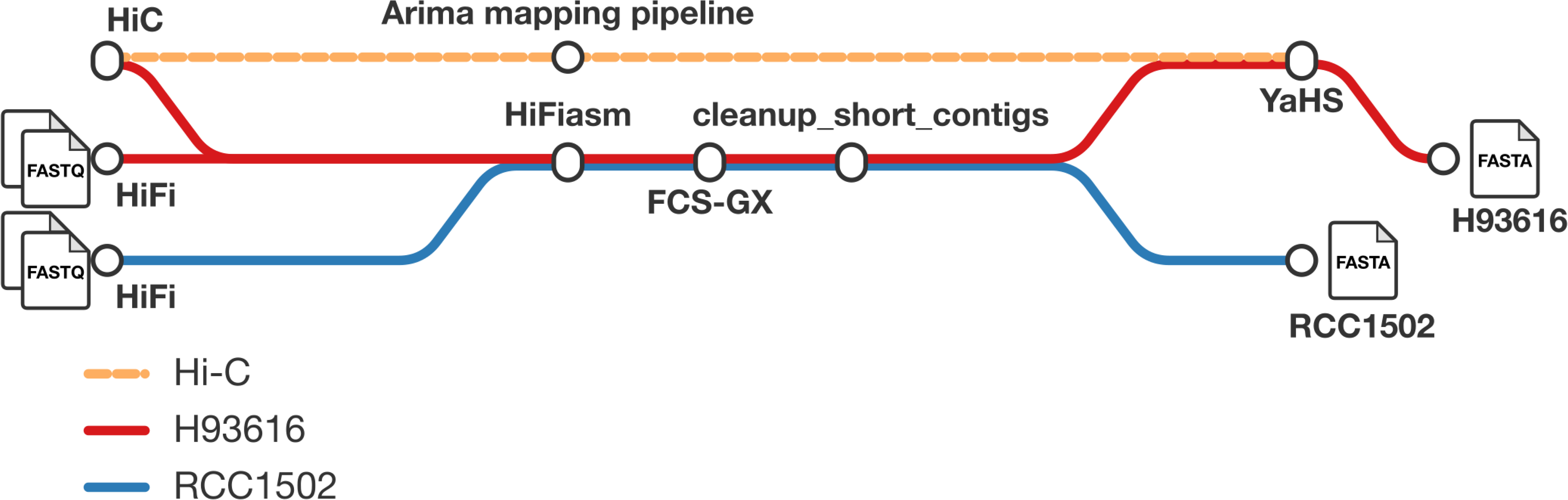
The workflow of *H. akashiwo* H93616 and RCC1502 genome assembly and scaffolding.

**Fig. 2.**
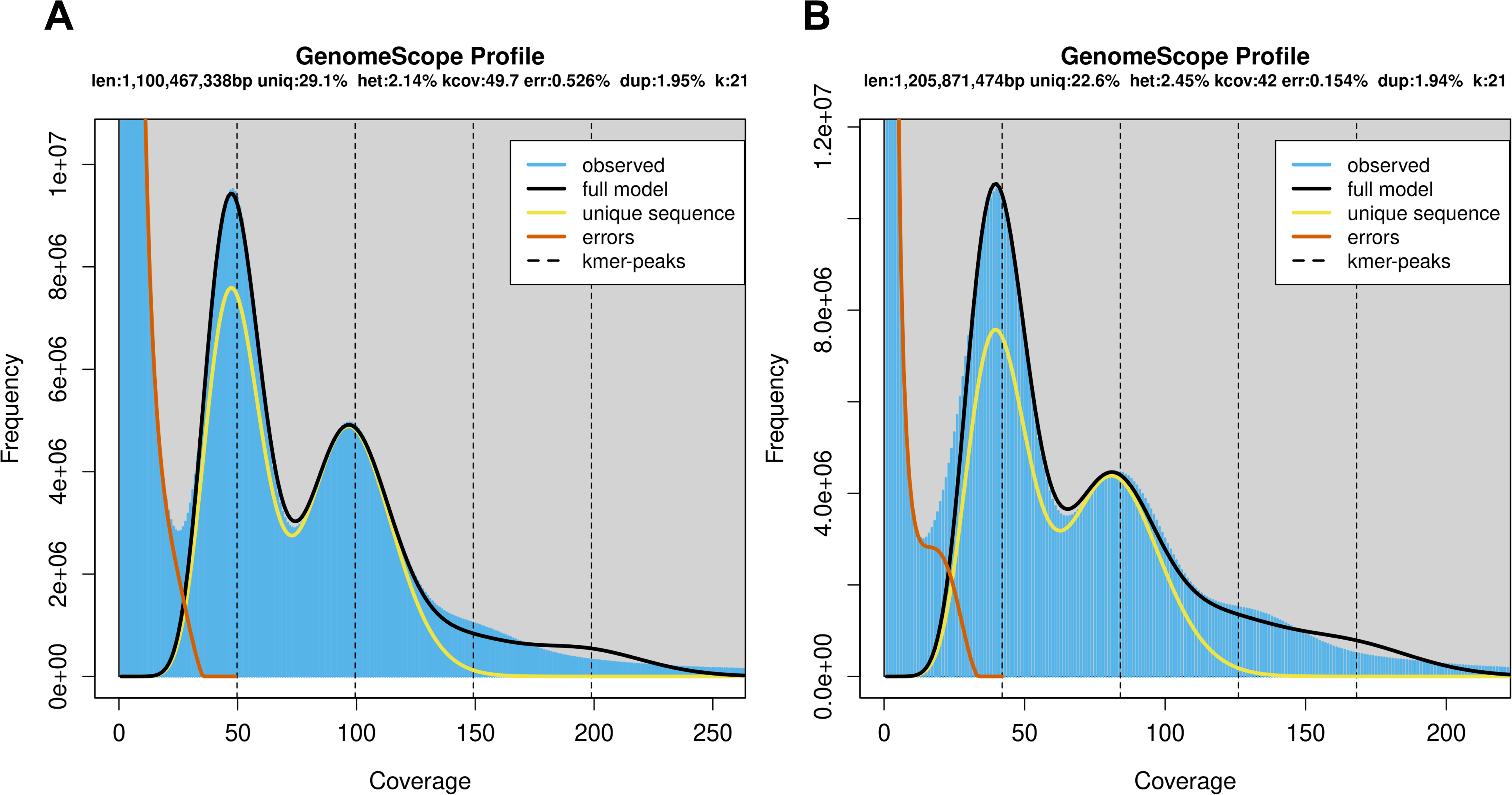
Estimation of genome sizes and heterozygosities of *H. akashiwo* H93616 (A) and RCC1502 (B).

**Table 1.** Statistics of sequence reads used for this study.

a) HiFi reads
| Strain | Total reads | Total bases (Gb) | Read N50 (bp) | >Q20 bases (Mb) | >Q20 rate (%) |
| --- | --- | --- | --- | --- | --- |
| H93616 | 4,739,923 | 86.5 | 18,203 | 73,432.1 | 85.4 |
| RCC1502 | 7,499,550 | 106.5 | 14,415 | 95,827.3 | 90.4 |

b) Hi-C reads and Illumina
| Before filtering |  |  |  |  | After filtering |  |  |  |
| --- | --- | --- | --- | --- | --- | --- | --- | --- |
| Strain | Total reads | Total bases (Gb) | >Q30 bases (Mb) | >Q30 rate (%) | Total reads | Total bases (Gb) | >Q30 bases (Mb) | >Q30 rate (%) |
| Hi-C |  |  |  |  |  |  |  |  |
| H93616 | 326,631,182 | 49.0 | 46,487.5 | 94.9 | 46.7 | 46,487.5 | 94.9 | 46.7 |
| Illumina PE |  |  |  |  |  |  |  |  |
| H93616 | 1,027,496,164 | 154.1 | 131,909.4 | 85.6 | 130.8 | 117,672.2 | 90.0 | 130.8 |
| RCC1502 | 877,881,288 | 131.7 | 127,512.2 | 96.8 | 130.1 | 126,462.4 | 97.2 | 130.1 |

### *De novo* assembly of *H. akashiwo* H93616 genome

The HiFi reads were assembled using hifiasm, producing a primary assembly with a total length of 1.3Gbp, comprising 2,418 contigs and an N_50_ of 31Mbp (Table 2). At this point, genome completeness, assessed by BUSCO genome mode, was as high as 91.0%, confirming the quality of the initial assembly.

**Table 2.** Assembly statistics of H93616 genome.

| Step | hifiasm | fcs-gx | cleaning | scaffolding | organella added |
| --- | --- | --- | --- | --- | --- |
| Number of contigs | 2,418 | 2,417 | 90 | 66 | 68 |
| Total length (base) | 1,308,794,501 | 1,303,925,422 | 1,179,221,920 | 1,179,226,620 | 1,179,443,489 |
| Max length (base) | 102,416,662 | 102,416,662 | 102,416,662 | 97,307,662 | 97,307,662 |
| N<sub>50</sub> | 31,170,977 | 31,170,977 | 34,926,071 | 65,983,599 | 65,983,599 |
| GC contents (%) | 45.71 | 45.71 | 47.24 | 47.24 | 47.24 |
| BUSCO complete<br>(%, genome mode) | 91.0 | 90.8 | 90.5 | 90.5 | 90.5 |

The primary contigs were screened for potential contamination using FCS-GX, leading to the removal of a single contig of bacterial origin. Contigs corresponding to mitochondrial and chloroplast genomes, rDNA clusters, and short self-overlapping sequences were further removed; this curation step yielded 90 contigs of nuclear origin, with a total length of 1.18 Gbp and an improved N50 of 34 Mbp. The GC content of the final contig set was 47.24% (Table 2), and the score calculated by BUSCO genome mode was 90.5% (Table 2, cleaning).

Subsequent scaffolding using Hi-C data and YaHS resulted in a subchromosome-scale assembly consisting of 66 scaffolds, with a total assembly length of 1.18Gbp and an N50 of 66Mbp.

Telomeric repeat sequences, AACCCT, were identified using the TIDK explore mode (Supplemental Fig 1). Eleven contigs were found to contain telomeric repeats at both ends, indicating putative complete chromosomal sequences, while an additional thirteen contigs harbored the repeats at one end.

The mitochondrial genome was identified from sequences tagged by cleanup_short_contigs. Contigs with lengths comparable to the previously reported mitochondrial genome (38,764 bp; accession KU561547) were aligned to the reference using MAFFT. Among these, a contig of 39,815 bp was selected as the mitochondrial genome based on sequence similarity.

Similarly, the chloroplast genome was identified from contigs removed by cleanup_short_contigs. Candidate sequences were aligned to a previously reported chloroplast reference genome (LC269923) using MAFFT and inverted repeats were identified by D-GENIES. The orientation of the candidate sequences was verified by alignment in both forward and reverse-complement directions. Redundant regions arising from the circular structure of the chloroplast genome were subsequently trimmed to obtain a non-redundant representation of the complete chloroplast sequence (Table 2, organelle added).

### Draft Genome Assembly of *H. akashiwo* strain RCC1502

For strain RCC1502, the HiFi reads were assembled using hifiasm, generating a primary assembly with a total length of 1.6 Gbp, comprising 4,732 contigs and an N50 of 28Mbp (Table 3). The GC content of this assembly was 45.40%, and genome completeness, assessed by BUSCO genome mode, was 90.7%.

**Table 3.** Assembly statistics of RCC1502 genome.

| Step | hifiasm | fcs-gx | cleaning |
| --- | --- | --- | --- |
| Number of contigs | 4,732 | 4,711 | 223 |
| Total length (base) | 1,612,051,257 | 1,611,191,762 | 1,430,577,794 |
| Max length (base) | 89,403,536 | 89,403,536 | 89,403,536 |
| N<sub>50</sub> | 27,964,728 | 29,385,481 | 33,446,378 |
| GC contents (%) | 45.40 | 45.40 | 47.26 |
| BUSCO complete<br>(%, genome mode) | 90.7 | 90.7 | 90.7 |

The resulting contigs were subsequently screened for potential contamination using FCS-GX, resulting in the removal of 21 contigs of bacterial origin. Further curation using cleanup_short_contigs yielded 223 contigs of nuclear origin, with a total length of 1.43 Gbps and an improved N50 of 33 Mbp. The GC content of the curated assembly was 47.26%, and a completeness score calculated by BUSCO genome mode was 90.7%.

Telomeric repeat sequences were identified in the curated dataset using the TIDK explore mode (Supplemental Table 1). Five contigs were found to contain telomeric repeats at both ends, suggesting putative complete chromosomal sequences, while an additional 26 contigs harbored telomeric repeats at one end.

### Gene prediction, and structural and functional annotations

Next, *H. akashiwo* genes were predicted from H93616 genome assembly. First, a tool for structural gene annotations based on a deep-learning approach, Helixer was adopted. About 42,500 genes were predicted when analyzed by land plant mode, while 16,360 genes were identified when analyzed by vertebrate mode; merging these results yielded 46,316 predicted genes. BUSCO scores obtained by protein mode for these results were as low as 13.1% ∼ 21.7%, indicating that these annotation approaches may not be suitable for the organism (Table 4).

**Table 4.**
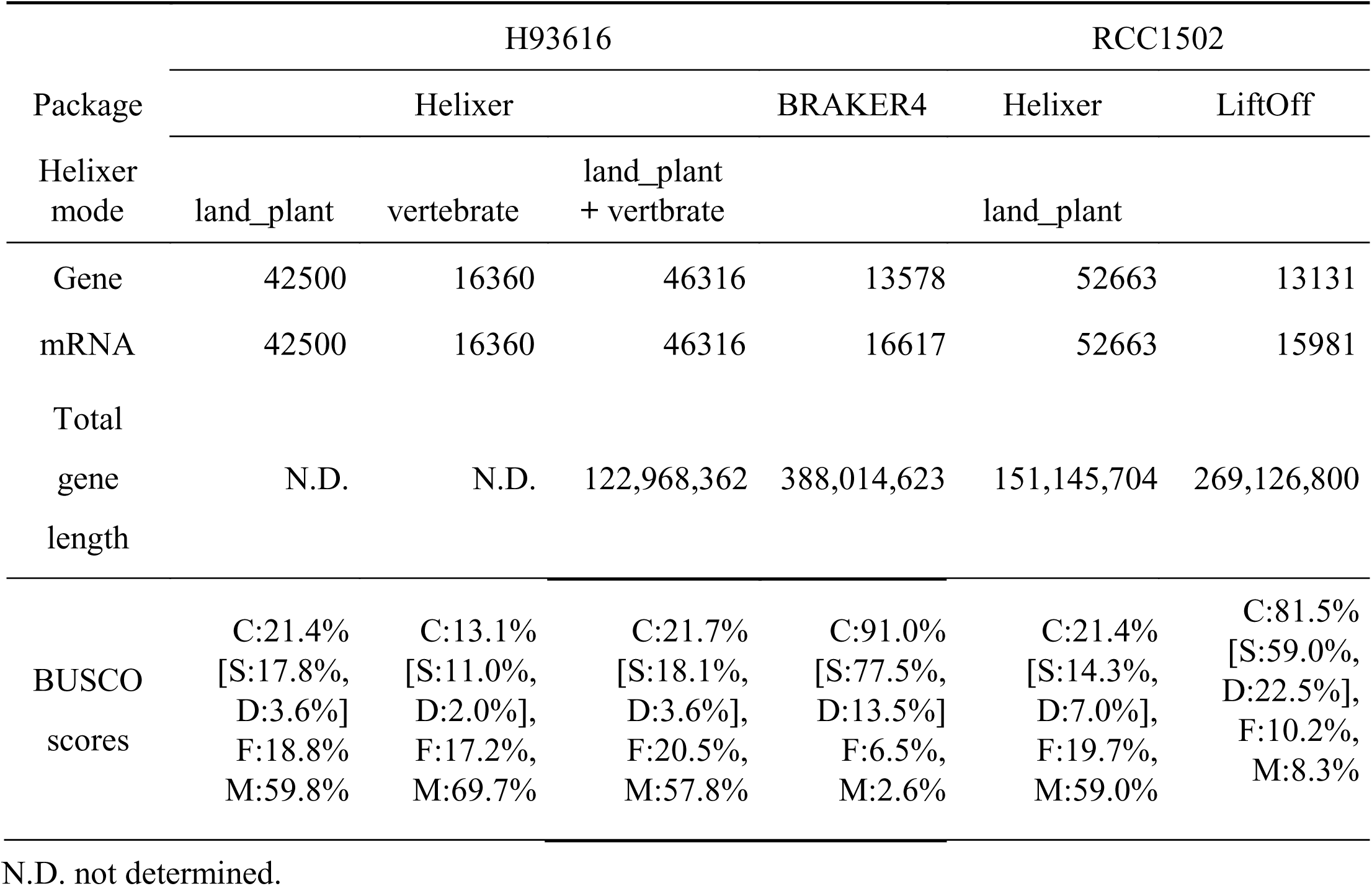
Gene prediction Genes encoded by genomes of strains H93616 and RCC5102 were predicted by various packages with different modes. C; complete, S; single, D; duplicate, F; fragmented, and M; missing.

Next, we adopted an evidence-based approach. H93616 genome assembly as well as RNA-seq reads previously published were analyzed by the package BRAKER4, an automated pipeline for structural annotation of eukaryotic genomes. While the number of the genes predicted by the approach was as little as 13,578, total gene length increased to 388 Mbp from 123 Mbp, which was obtained by Helixer (Table 4). In addition, BUSCO complete score by protein mode was elevated to 91%. These results indicate that the BRAKER4 gave the best prediction for genes coded by H93616 among the approaches tested here (Table 4).

For gene prediction for RCC1502, Helixer and Liftoff were adopted. About 52,600 genes were predicted by Helixer run with land plant mode, while 13,131 genes were identified by Liftoff (Table 4). Total gene length based on Liftoff was calculated to be 269 Mbp, compare to 151 Mbp obtained based on Helixer prediction (Table 4). BUSCO completeness scores obtained by protein mode were 21.4% for Helixer-based prediction and 81.5% for Liftoff-based prediction, respectively (Table 4).

Genes coded by H93616 genome were further annotated for their protein structure and function. Among ∼13,000 total predicted genes, 4447 genes were predicted by KAAS for their functions, linking to various cellular and metabolic pathways. In addition, 1104 genes were predicted to possesses characteristic protein domains and structure. Both function and protein structure annotations are provided as Supplemental data.

### Synteny between H93616 and RCC1502 genomes

Finally, synteny between H93616 and RCC1502 genome assemblies was analyzed by GENESPACE (Fig 3). Overall, synteny between the two strains are widely conserved. Several short RCC1502 scaffolds aligned to long scaffolds of H93616, such as scaffolds 1 and 3, likely suggesting that they are part of the corresponding chromosomes. On the other hand, part of the RCC1502 scaffolds, such as the ones in ptg000033I or ptg150000I, were apparently aligned to the parts of separate H93616 contigs, suggesting that these segments putatively translocated. In addition, some segments of scaffolds of either strain are aligned to separate scaffolds of the other strain more than two times, suggesting that these parts may have duplicated.

**Fig. 3.**
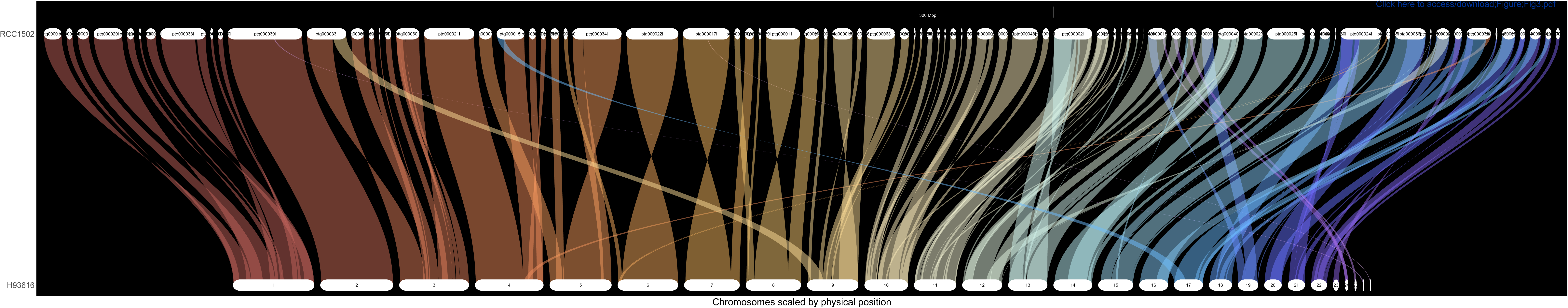
Genome synteny between H93616 (bottom) and RCC1502 (top). The H93616 scaffolds are arranged in order of their length. The scaffolds/contigs that do not have syntenic counterparts are not shown in the figure.

## Discussion

*H. akashiwo* belongs to the class Raphidophyceae, division Ochrophyta within the stramenopile lineage. As of 30 June 2026, the NCBI Genome database contained complete or chromosome-scale genome assemblies for 38 ochrophyte species, with genome sizes ranging up to 1.09 Gb in *Epithemia catenata* (GCA_964019935.2, see Supplemental Table 2). Only a limited number of these genomes have been comprehensively annotated. In the present study, we report high-quality genome assemblies for two *H. akashiwo* strains, H93616 and RCC1502. Because long-read sequence datasets were subjected to assembly, the length and quality of assemblies were dramatically improved compared to the currently available, scaffold-level *H. akashiwo* genome assemblies in the database. The assembly of strain H93616 comprises 66 scaffolds, including eleven scaffolds capped by canonical telomeric repeats at both termini, and exhibits an N50 of approximately 65.9 Mb, indicative of a subchromosomal-level assembly. The assembly of strain RCC1502 consists of 223 scaffolds, with five contigs containing telomeric repeats at both ends. Given the estimated genome size of approximately 1.1 Gb based on *k-*mer analysis, these assemblies rank among the largest and most contiguous genomes currently available for Stramenopiles Ochrophyta, and only available genome sequences for Raphidophyceae. Together, our study substantially expands the genomic resources for the taxa.

For structural genome annotation, we initially adopted Helixer, a widely used gene prediction framework that combines homology-based inference with deep learning. However, the available land plant and invertebrate models were not suitable for *H. akashiwo*: BUSCO v6 analysis in protein mode showed that the predicted gene sets recovered only approximately 20% of stramenopile core genes while genome sequence completeness predicted by the same package run by genome mode exceed 90%. We therefore adopted an evidence-based gene prediction approach using BRAKER4, incorporating RNA-seq data generated from strain H93616. These RNA-seq data represent a subset of the sequencing reads previously used for transcriptome shotgun assembly (TSA; accession ICRV01, [63]) published by our group.

The BRAKER4-based annotation yielded substantially improved results relative to the Helixer predictions. BUSCO v6 completeness assessed in protein mode increased to approximately 90%, with both the average coding sequence (CDS) length and the number of predicted exons per gene being markedly increased, indicating improved gene model completeness and structural accuracy. The incorporation of additional RNA-seq datasets obtained under a broader range of physiological conditions may further improve genome annotation. For example, we recently demonstrated that *H. akashiwo* can utilize bacterial polyphosphate as a phosphorus source through a heterotrophic nutritional strategy involving bacterivory and digestion [31].

Because the RNA-seq data used in the present study were generated under conditions in which *H. akashiwo* primarily relied on photosynthesis, genes associated with bacterivory and other heterotrophic processes may not have been actively expressed. Incorporating transcriptomic datasets derived from conditions that induce heterotrophic feeding may improve the identification and annotation of additional genes in the genome.

Credibility of the gene prediction is further supported by the protein-level annotation; More than 80% of the *H. akashiwo* gene products were associated with one or more signature protein motif, and ∼ 33% were characterized to be homologous to KEGG genes that are defined with functions at significant level. Interestingly, Helixer gene prediction using the land-plant model produced highly variable results across Ochrophyta genomes, ranging between 20.9% to 94.3%, when assessed with BUSCO v6 in protein mode (Supplemental Table 2). Of the 38 Ochrophyta species examined, 16 achieved >80% BUSCO completeness when analyzed by protein mode, whereas three species exhibited completeness scores below 25%, likewise the two *H. akashiwo* strains analyzed in this study, which each scored approximately 20%. This substantial variation suggests that some stramenopile lineages may possess genomic features that are not adequately captured by the Helixer models currently available.

One possible explanation is the presence of atypical RNA splicing mechanisms in the organism. For example, during BRAKER4-based annotation of *H. akashiwo*, we identified several coding sequences supported by transcriptomic evidence that appeared to utilize splice sites inconsistent with the canonical GU-AG splice-site rule. These observations raise the possibility that *H. akashiwo* and some other taxa employ non-canonical splicing patterns that may compromise the performance of gene prediction models trained predominantly on conventional eukaryotic splice-site architectures. Rigorous evidence-based annotation of the *H. akashiwo* genome, coupled with a detailed characterization of its splicing mechanisms, may improve gene prediction accuracy for this species and facilitate the development of more robust annotation strategies for other ochrophyte and stramenopile genomes exhibiting similarly low prediction performance.

Genome synteny analysis revealed that overall synteny was well conserved between H93616 and RCC1502, that are originated in distant locations. On the other hand, there are some partial inconsonance in some corresponding scaffolds, that may imply the major structural rearrangements, large-scale duplications or simply miss-assembly of either genome. Incorporation of longer reads, such as Oxford Nanopore sequencing, or Hi-C reads to RCC1502 assembly may clarify this point.

We previously reported that the sequences of two open reading frames (ORFs) encoded by the mitochondrial genomes of *H. akashiwo* strains from diverse geographic locations exhibited a clear pattern of isolation by distance [18, 19, 64]. Notably, one of these genes, designated *mtORFvar1*, displayed marked geographic variation. Strains isolated from regions north of 42°N, represented by RCC1502, possess an extended N-terminal sequence that is absent in strains from lower-latitude regions, such as H93616 [18, 19, 64].

Comparative analyses of the nuclear genomes and gene repertoires of RCC1502 and H93616, and the identification and characterization of genes with variations, may provide valuable insights into the mechanisms underlying regional adaptation in *H. akashiwo* and help elucidate the species’ dispersal history.

In this study, we produced high-quality genome assemblies and comprehensive gene annotations for two *H. akashiwo* strains, thereby providing the first extensive genomic resources for this species. These resources establish a valuable foundation for future studies of the physiology, environmental adaptation, evolution, and particularly, bloom-forming mechanisms of *H. akashiwo*.

## Acknowledgements

We thank the technical staff at the National Institute of Genetics for their assistance with NGS sequencing. This work was supported by JSPS KAKENHI Grant Number JP221S0002, JP16H06279, 24K21840, 26K22969 and “Initiative for Realizing Diversity in the Research Environment” from MEXT, Japan. We sincerely thank the Nagase Science and Technology Foundation and the Casio Science Foundation for financially supporting this research on *H. akashiwo* since its inception.

## Data Availabilities

The raw sequencing reads generated in this study have been deposited in the DDBJ Sequence Read Archive (DRA) under BioProject accession PRJDB35616 (DRA accession numbers DRR1079267 - DRR1079269) for H93616 and PRJDB42657 (DRA accession numbers DRR1079270 - DRR1079272) for RCC1502. The genome assembly of H93616 has been deposited in DDBJ under accession BAALQZ010000001-BAALQZ010000066, LC941944 (mitochondrion genome), and LC941945 (chloroplast genome). The genome assembly of RCC1502 has been deposited in DDBJ under accession BAALQW010000001-BAALQW010000223.

## Notes

### Competing Interest Statement

The authors have declared no competing interest.

## References

1. Hallegraeff GM. A review of harmful algal blooms and their apparant global increase. Phycologia 1993;1993:79–99. 10.2216/i0031-8884-32-2-79.1

2. Anderson DM, Glibert PM, Burkholder JM. Harmful algal blooms and eutrophication: Nutrient sources, composition, and consequences. Estuaries 2002;25:704–25. 10.1007/BF02804901

3. Backer LC, McGillicuddy DJ. Harmful algal blooms: At the interface between coastal oceanography and human health. Oceanography 2006;19:96–106. 10.5670/oceanog.2006.72

4. Fu FX, Tatters AO, Hutchins DA. Global change and the future of harmful algal blooms in the ocean. Marine Ecology Progress Series 2012;470:207–33. 10.3354/Meps10047

5. Maso M, Garces E. Harmful microalgae blooms (hab); problematic and conditions that induce them. Mar Pollut Bull 2006;53:620–30. 10.1016/j.marpolbul.2006.08.006

6. Heisler J, Glibert PM, Burkholder JM, Anderson DM, Cochlan W, Dennison WC et al. Eutrophication and harmful algal blooms: A scientific consensus. Harmful Algae 2008;8:3–13. 10.1016/j.hal.2008.08.006

7. Smith VH, Schindler DW. Eutrophication science: Where do we go from here? Trends Ecol Evol 2009;24:201–7. 10.1016/j.tree.2008.11.009

8. Eppley R Temperature and phytoplankton growth in the sea. Fish Bull. 1063–85.

9. Rensel JEJ, Haigh N, Tynan TJ. Fraser river sockeye salmon marine survival decline and harmful blooms of heterosigrna akashiwo. Harmful Algae 2010;10:98–115. 10.1016/j.hal.2010.07.005

10. Smayda TJ Ecophysiology and bloom dynamics of heterosgma akashiwo (raphidophyceae). In: Anderson DM, Cembella AD, Hallegraeff GF (eds.), Physiological ecology of harmful algal blooms, New York: Springer-Verlag. 113–31.

11. Haque SM, Onoue Y. Variation in toxin compositions of two harmful raphidophytes, chattonella antiqua and chattonella marina, at different salinities. Environmental Toxicology 2002;17:113–18. 10.1002/Tox.10039

12. Haque SM, Onoue Y. Effects of salinity on growth and toxin production of a noxious phytoflagellate, heterosigma akashiwo (raphidophyceae). Botanica Marina 2002;45:356–63. 10.1515/Bot.2002.036

13. Mohamed ZA, Al-Shehri AM. The link between shrimp farm runoff and blooms of toxic heterosigma akashiwo in red sea coastal waters. Oceanologia 2012;54:287–309. 10.5697/Oc.54-2.287

14. Yang CZ, Albright LJ, Yousif AN. Oxygen-radical-mediated effects of the toxic phytoplankter heterosigma carterae on juvenile rainbow-trout oncorhynchus mykiss. Diseases of Aquatic Organisms 1995;23:101–08. 10.3354/Dao023101

15. Kempton J, Keppler CJ, Lewitus A, Shuler A, Wilde S. A novel heterosigma akashiwo (raphidophyceae) bloom extending from a south carolina bay to offshore waters. Harmful Algae 2008;7:235–40. 10.1016/j.ha1.2007.08.003

16. Khan S, Arakawa O, Onoue Y. Neurotoxins in a toxic red tide of heterosigma akashiwo (raphidophyceae) in kagoshima bay, japan. Aquaculture Research 1997;28:9–14. 10.1111/j.1365-2109.1997.tb01309.x

17. Ono K, Khan S, Onoue Y. Effects of temperature and light intensity on the growth and toxicity of heterosigma akashiwo (raphidophyceae). Aquaculture Research 2000;31:427–33. 10.1046/j.1365-2109.2000.00463.x

18. Ueki S. Phylogeographic characteristics of hypervariable regions in the mitochondrial genome of a cosmopolitan, bloom-forming raphidophyte, heterosigma akashiwo. J Phycol 2019;55:858–67. 10.1111/jpy.12868

19. Higashi A, Nagai S, Salomon PS, Ueki S. A unique, highly variable mitochondrial gene with coding capacity of heterosigma akashiwo, class raphidophyceae. Journal of Applied Phycology 2017 10.1007/s10811-017-1142-2

20. Engesmo A, Eikrem W, Seoane S, Smith K, Edvardsen B, Hofgaard A et al. New insights into the morphology and phylogeny ofheterosigma akashiwo(raphidophyceae), with the description of heterosigma minor sp. Nov. Phycologia 2016;55:279–94. 10.2216/15-115.1

21. Martinez R, Orive E, Laza-Martinez A, Seoane S. Growth response of six strains of heterosigma akashiwo to varying temperature, salinity and irradiance conditions. Journal of Plankton Research 2010;32:529–38. 10.1093/plankt/fbp135

22. Shikata T, Nagasoe S, Matsubara T, Yamasaki Y, Shimasaki Y, Oshima Y et al. Effects of temperature and light on cyst germination and germinated cell survival of the noxious raphidophyte heterosigma akashiwo. Harmful Algae 2007;6:700–06. 10.1016/j.hal.2007.02.008

23. Fu FX, Zhang YH, Warner ME, Feng YY, Sun J, Hutchins DA. A comparison of future increased co2 and temperature effects on sympatric heterosigma akashiwo and prorocentrum minimum. Harmful Algae 2008;7:76–90. 10.1016/j.hal.2007.05.006

24. Xu D, Zhou B, Wang Y, Ju Q, Yu QY, Tang XX. Effect of co2 enrichment on competition between skeletonema costatum and heterosigma akashiwo. Chinese Journal of Oceanology and Limnology 2010;28:933–39. 10.1007/s00343-010-9071-9

25. Han MS, Kim YP, Cattolico RA. Heterosigma akashiwo (raphidophyceae) resting cell formation in batch culture: Strain identity versus physiological response. Journal of Phycology 2002;38:304–17. 10.1046/j.1529-8817.2002.01087.x

26. Imai I, Itakura S. Importance of cysts in the population dynamics of the red tide flagellate heterosigma akashiwo (raphidophyceae). Mar Biol 1999;133:755–62. 10.1007/s002270050517

27. Sengupta A, Carrara F, Stocker R. Phytoplankton can actively diversify their migration strategy in response to turbulent cues. Nature 2017;543:555–58. 10.1038/nature21415

28. Jeong HJ. Mixotrophy in red tide algae raphidophytes. Journal of Eukaryotic Microbiology 2011;58:215–22. 10.1111/j.1550-7408.2011.00550.x

29. Jeong HJ, Seong KA, Du Yoo Y, Kim TH, Kang NS, Kim S et al. Feeding and grazing impact by small marine heterotrophic dinoflagellates on heterotrophic bacteria. Journal of Eukaryotic Microbiology 2008;55:271–88. 10.1111/j.1550-7408.2008.00336.x

30. Seong KA, Jeong HJ, Kim S, Kim GH, Kang JH. Bacterivory by co-occurring red-tide algae, heterotrophic nanoflagellates, and ciliates. Marine Ecology Progress Series 2006;322:85–97. 10.3354/Meps322085

31. Fukuyama S, Usami F, Hirota R, Satoh A, Ohara S, Kondo K et al. Proliferation of a bloom-forming phytoplankton via uptake of polyphosphate-accumulating bacteria under phosphate-limiting conditions. ISME Commun 2025;5:ycaf192. 10.1093/ismeco/ycaf192

32. Tsukahara S, Bousios A, Perez-Roman E, Yamaguchi S, Leduque B, Nakano A et al. Centrophilic retrotransposon integration via cenh3 chromatin in arabidopsis. Nature 2025;637:744–48. 10.1038/s41586-024-08319-7

33. Kokot K, Długosz M, Deorowicz S. Kmc 3: Counting and manipulating k-mer statistics. Bioinformatics 2017;33:2759–61. 10.1093/bioinformatics/btx304

34. Vurture GW, Sedlazeck FJ, Nattestad M, Underwood CJ, Fang H, Gurtowski J et al. Genomescope: Fast reference-free genome profiling from short reads. Bioinformatics 2017;33:2202–04. 10.1093/bioinformatics/btx153

35. Chen S. Fastp 1.0: An ultra-fast all-round tool for fastq data quality control and preprocessing. iMeta 2025;2:e107. 10.1002/imt2.107

36. Chen S. Ultrafast one-pass fastq data preprocessing, quality control, and deduplication using fastp. iMeta 2023;2:e107. 10.1002/imt2.107

37. Chen S, Zhou Y, Chen Y, Gu J. Fastp: An ultra-fast all-in-one fastq preprocessor. Bioinformatics 2018;34:i884–i90. 10.1093/bioinformatics/bty560

38. Cheng H, Concepcion GT, Feng X, Zhang H, H. L. Haplotype-resolved de novo assembly using phased assembly graphs with hifiasm. Nat Methods 2021;18:170–75. 10.1038/s41592-020-01056-5

39. Cheng H, Jarvis ED, Fedrigo O, Koepfli KP, Urban L, Gemmell NJ, et al. Haplotype-resolved assembly of diploid genomes without parental data. Nat Biotechnol 2022;40:1332–35. 10.1038/s41587-022-01261-x

40. Cheng H, Asri M, Lucas J, Koren S, Li H. Scalable telomere-to-telomere assembly for diploid and polyploid genomes with double graph. Nat Methods 2024;21:967–70. 10.1038/s41592-024-02269-8

41. Astashyn A, Tvedte ES, Sweeney D, Sapojnikov V, Bouk N, Joukov V et al. Rapid and sensitive detection of genome contamination at scale with fcs-gx. Genome Biol 2024;25:60. 10.1186/s13059-024-03198-7

42. Shen W, Sipos B, Zhao L. Seqkit2: A swiss army knife for sequence and alignment processing. iMeta 2024;3:e191. 10.1002/imt2.191

43. Manni M, Berkeley M, Seppey M, Simão FA, Zdobnov EM. Busco update: Novel and streamlined workflows along with broader and deeper phylogenetic coverage for scoring of eukaryotic, prokaryotic, and viral genomes. Molecular Biology and Evolution 2021;38:4647–54. 10.1093/molbev/msab199

44. Tegenfeldt F, Kuznetsov D, Manni M, Berkeley M, Zdobnov EM, Kriventseva EV. Orthodb and busco update: Annotation of orthologs with wider sampling of genomes. Nucleic Acids Research 2025;53:D516–D22. 10.1093/nar/gkae987

45. Zhou C, McCarthy SA, Durbin R. Yahs: Yet another hi-c scaffolding tool. Bioinformatics;39:btac808. 10.1093/bioinformatics/btac808

46. Durand NC, Robinson JT, Shamim MS, Machol I, ¥., Mesirov JP, Lander ES et al. Juicebox provides a visualization system for hi-c contact maps with unlimited zoom. Cell Systems 2016;3:99–101. 10.1016/j.cels.2015.07.012

47. Brown MR, de La Rosa PMG, Blaxter M. Tidk: A toolkit to rapidly identify telomeric repeats from genomic datasets. Bioinformatics 2025;41:btaf049. 10.1093/bioinformatics/btaf049

48. Katoh K, Standley DM. Mafft multiple sequence alignment software version 7: Improvements in performance and usability. Molecular Biology and Evolution 2013;30:772–80. 10.1093/molbev/mst010

49. Katoh K, Standley DM. A simple method to control over-alignment in the mafft multiple sequence alignment program. Bioinformatics 2016;32:1933–42. 10.1093/bioinformatics/btw108

50. Cabanettes F, Klopp C. D-genies: Dot plot large genomes in an interactive, efficient and simple way. Peer J 2018;6:e4958. 10.7717/peerj.4958

51. Holst F, Bolger AM, Kindel F, Günther C, Maß J, Triesch S et al. Helixer: Ab initio prediction of primary eukaryotic gene models combining deep learning and a hidden markov model. Nat Methods 2025;23:732–39. 10.1038/s41592-025-02939-1

52. Stiehler F, Steinborn M, Scholz, Dey D, Weber APM, Denton AK. Helixer: Cross-species gene annotation of large eukaryotic genomes using deep learning. Bioinformatics 2020:btaa1044. 10.1093/bioinformatics/btaa1044

53. Stanke M, Diekhans M, Baertsch R, Haussler D. Using native and syntenically mapped cdna alignments to improve de novo gene finding. Bioinformatics 2008;24:637–44. 10.1093/bioinformatics/btn013

54. M. S, Schöffmann O, Morgenstern B. Gene prediction in eukaryotes with a generalized hidden markov model that uses hints from external sources. BMC Bioinformatics 2006;7:62. 10.1186/1471-2105-7-62

55. Gabriel L, Bruna T, Hoff KJ, Borodovsky M, Stanke M. Tsebra: Transcript selector for braker. BMC Bioinformatics 2021;22:566. 10.1186/s12859-021-04482-0

56. Gabriel L, Bruna T, Hoff KJ, Ebel M, Lomsadze A, Borodovsky M et al. Braker3: Fully automated genome annotation using rna-seq and protein evidence with genemark-etp, augustus and tsebra. Genome Res 2024;34:769–77. 10.1101/gr.278090.123

57. Brůna T, Lomsadze A, Borodovsky M. Genemark-etp significantly improves the accuracy of automatic annotation of large eukaryotic genomes. Genome Res 2024;34:757–68. 10.1101/gr.278373.123

58. Kovaka S, Zimin AV, Pertea GM, Razaghi R, Salzberg SL, Pertea M. Transcriptome assembly from long-read rna-seq alignments with stringtie2. Genome Biol 2019;20:1–13. 10.1186/s13059-019-1910-1

59. Shumate A, Salzberg SL. Liftoff: Accurate mapping of gene annotations. Bioinformatics 2021;37:1639–43. 10.1093/bioinformatics/btaa1016

60. Moriya Y, Itoh M, Okuda S, Yoshizawa AC, Kanehisa M. Kaas: An automatic genome annotation and pathway reconstruction server. Nucleic Acids Research 2007;35:W182–W85. 10.1093/nar/gkm321

61. Jones P, Binns D, Chang H-Y, Fraser M, Li W, McAnulla C et al. Interproscan 5: Genome-scale protein function classification. Bioinformatics 2014;30:1236–40. 10.1093/bioinformatics/btu031

62. Lovell JT, Sreedasyam A, Schranz EM, Wilson M, Carlson JW, Harkess A et al. Genespace tracks regions of interest and gene copy number variation across multiple genomes. eLife 2022;11:e78526. 10.7554/eLife.78526

63. Sato M, Seki M, Suzuki Y, Ueki S. The dataset of de novo assembly and inferred functional annotation of the transcriptome of heterosigma akashiwo äi0, a bloom-forming, cosmopolitan raphidophyte Data in Brief 2023;48:109071. 10.1016/j.dib.2023.109071

64. Higashi A, Nagai S, Seone S, Ueki S. A hypervariable mitochondrial protein coding sequence associated with geographical origin in a cosmopolitan bloom-forming alga, heterosigma akashiwo. Biol Lett 2017;13 10.1098/rsbl.2016.0976

