## Supplementary figures and images for "Whole genome sequences and annotations of Japanese and French strains of *Heterosigma akashiwo*"

### Supplemental Fig 1

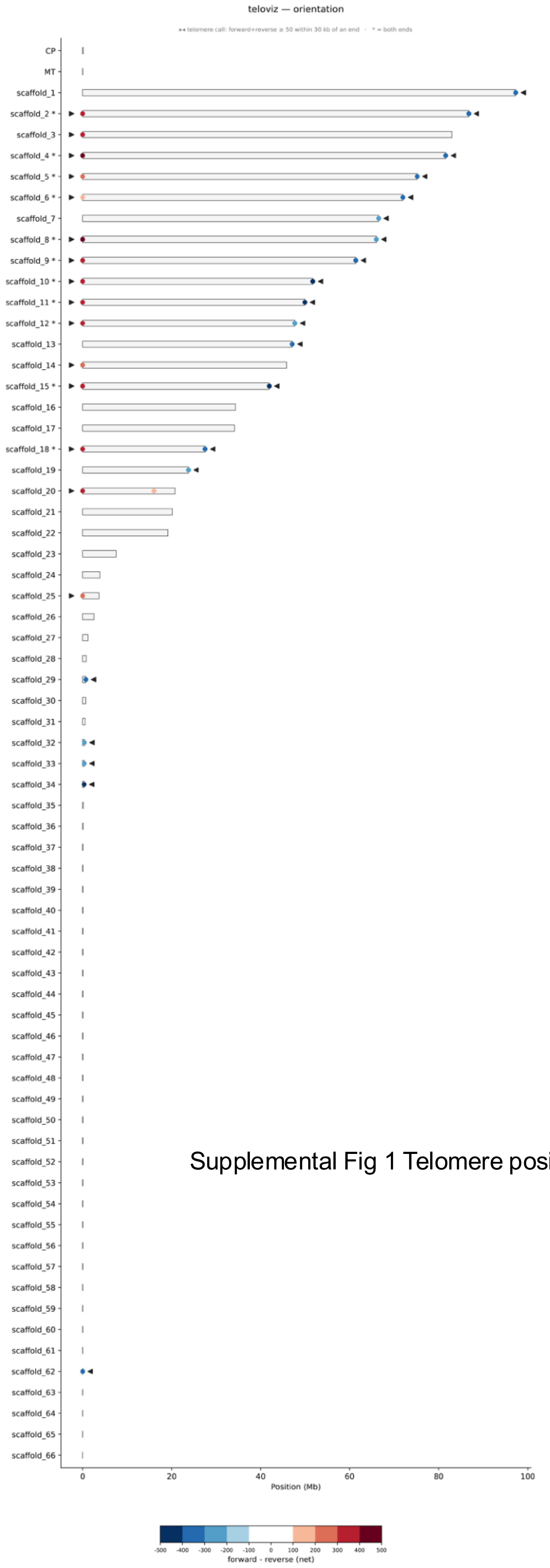


Supplemental Fig 1 Telomere positions on H93616 scaffolds
